# NEAT-seq: Simultaneous profiling of intra-nuclear proteins, chromatin accessibility, and gene expression in single cells

**DOI:** 10.1101/2021.07.29.454078

**Authors:** Amy F Chen, Benjamin Parks, Arwa S Kathiria, Benjamin Ober-Reynolds, Jorg Goronzy, William J Greenleaf

## Abstract

Oligonucleotide-conjugated antibodies^1^ have allowed for joint measurement of surface protein abundance and the transcriptome in single cells using high-throughput sequencing. Extending these measurements to gene regulatory proteins in the nucleus would provide a powerful means to link changes in abundance of trans-acting TFs to changes in activity of cis-acting elements and expression of target genes. Here, we introduce Nuclear protein Epitope, chromatin Accessibility, and Transcriptome sequencing (NEAT-seq), a technique to simultaneously measure nuclear protein abundance, chromatin accessibility, and the transcriptome in single cells. We apply this technique to profile CD4 memory T cells using a panel of master transcription factors (TFs) that drive distinct helper T cell subsets and regulatory T cells (Tregs) and identify examples of TFs with regulatory activity gated by three distinct mechanisms: transcription, translation, and regulation of chromatin binding. Furthermore, we identify regulatory elements and target genes associated with each TF, which we use to link a non-coding GWAS SNP within a GATA motif to both strong allele-specific chromatin accessibility in cells expressing high levels of GATA3 protein, and a putative target gene.

## Main

Recent progress in diverse multimodal single cell technologies has revolutionized our ability to characterize cell states and identify gene regulatory programs across various cell types. For example, methods pairing ATAC-seq and RNA-seq in single cells have allowed association of epigenetic status with transcriptional output, enabling identification of putative target genes of regulatory elements and uncovering regulatory mechanisms controlling specific gene networks, such as epigenetic priming of genes during development^2^. Antibodies linked to barcoded oligonucleotides have enabled surface protein measurements using a sequencing read-out, which, when combined with RNA- or ATAC-seq in single cells, has been particularly informative for profiling immune populations traditionally isolated based on surface protein markers^1,3^. Recently, this approach was extended to measuring intracellular proteins^4–8^. However, quantification of nuclear gene regulatory proteins along with chromatin accessibility profiling and RNA-seq has not been demonstrated.

In particular, existing methods for linking single cell chromatin accessibility to transcription do not directly quantify the central driver of gene regulation: transcription factors. TFs bind directly to enhancer elements to drive target gene expression^9^, and therefore measuring their abundance is a critical factor in understanding gene regulatory processes. However, TF abundance is difficult to estimate via scRNA-seq due to the combination of low transcript abundance and relatively low capture rates^10,11^. Directly measuring single-cell protein levels of TFs, which are often much more abundant than their encoding transcripts^12^, can link individual TFs to regulated enhancers and target genes by correlating changes in TF protein levels to changes in regulatory element accessibility and expression of nearby genes. Such analyses incorporating TF protein measurements can help distinguish direct target genes from secondary effects, reveal cooperative and antagonistic effects of multiple TFs on gene regulation, and enable more accurate identification of TF-mediated gene regulatory networks that drive cell fate.

Here, we develop NEAT-seq, a method that enables sensitive and specific quantification of nuclear proteins along with ATAC-seq and RNA-seq in single cells. We use NEAT-seq to profile CD4 memory T cells, a functionally diverse cell population regulated by several known master TFs. We illustrate the utility of NEAT-seq for interrogating the relationship between master TF abundance, chromatin accessibility changes, and gene expression, identifying regulatory elements and candidate target genes associated with TFs driving distinct CD4 T cell lineages and uncovering three distinct regulatory mechanisms controlling expression and activity of the master TFs themselves. Using these data, we also nominate a candidate target gene for a GWAS SNP associated with pulmonary disease that lies within a GATA motif, and show that the SNP alters chromatin accessibility in a GATA3 dosage-dependent manner.

While multiple groups have demonstrated sequencing-based surface protein quantification using barcoded antibodies^1,3,13,14^, application to nuclear proteins has been challenging due to high levels of background oligo-antibody staining in the nucleus, potentially driven by the highly negatively charged conjugated single stranded DNA (ssDNA) oligo^5,15^. One approach to reduce non-specific staining is to saturate cells with single stranded nucleic acids or other negatively charged polymers in an attempt to block cellular components that bind non-specifically to ssDNA^5,6,8,15^. However, we hypothesized that directly blocking the charge of the antibody oligo might dramatically improve signal over background. We tested this hypothesis by binding the conjugated ssDNA oligo with E.coli single-stranded DNA binding protein (EcoSSB). Several features of EcoSSB made it an attractive candidate for this application. First, EcoSSB binds with high affinity to ssDNA in a sequence non-specific manner, but has low affinity for dsDNA in the nucleus^16^, thereby minimizing background by neutralizing oligonucleotide charge while likely not influencing global chromatin accessibility within the nucleus. Second, EcoSSB has a slow dissociation rate,^16,17^ enhancing its ability to remain complexed with the conjugated oligo during staining. Finally, since one of the biological functions of EcoSSB in cells is to facilitate DNA replication^18^, we anticipated that EcoSSB bound to the conjugated oligo would not interfere with PCR amplification of the oligo during library generation.

To test and optimize staining with EcoSSB-bound oligo-conjugated antibodies, we designed a system to easily and accurately assess the specificity and dynamic range of antibody signal produced under diverse staining conditions. We first conjugated an anti-GFP antibody with streptavidin and mixed the conjugated antibody with 80-bp ssDNA oligos modified with 5’ biotin and 3’ Cy5 fluorophore. This antibody allowed us to compare oligo-antibody staining levels (via Cy5 fluorescence) to GFP fluorescence levels within a cell. We used this antibody to stain HEK293 cells expressing a nuclear-localized GFP and measured GFP and Cy5 fluorescence using flow cytometry.

To test the effects of EcoSSB on the specificity of GFP staining using an oligo-antibody, we incubated nuclei expressing GFP with oligo-antibody that had been pre-incubated with or without EcoSSB. Using flow cytometry, we observed that pre-incubating the oligo-antibody with EcoSSB greatly reduced background staining and yielded a strong correlation between Cy5 antibody signal and GFP levels within the nucleus that was not observed in the absence of EcoSSB (Fig. 1a). To confirm that our quantification of conjugated oligo reflected GFP protein levels within the nucleus, we sorted nuclei into 3 populations of increasing GFP expression for qPCR targeting the conjugated oligonucleotide (Supplementary Fig. 1a). The relative levels of conjugated oligo quantified from each population via qPCR corresponded to the differences in GFP abundance measured through flow cytometry (Supplementary Fig. 1b). To determine if these staining conditions would be sufficiently sensitive to detect endogenous levels of nuclear protein, we stained for the TF GATA1 in K562 cells, using embryonic stem cells (ESCs) as a negative control. Indeed, we observed a marked increase in GATA1 staining in K562 relative to ESCs using a GATA1 antibody linked to the Cy5-modified oligo (Supplementary Fig. 1c).

**Fig. 1:**
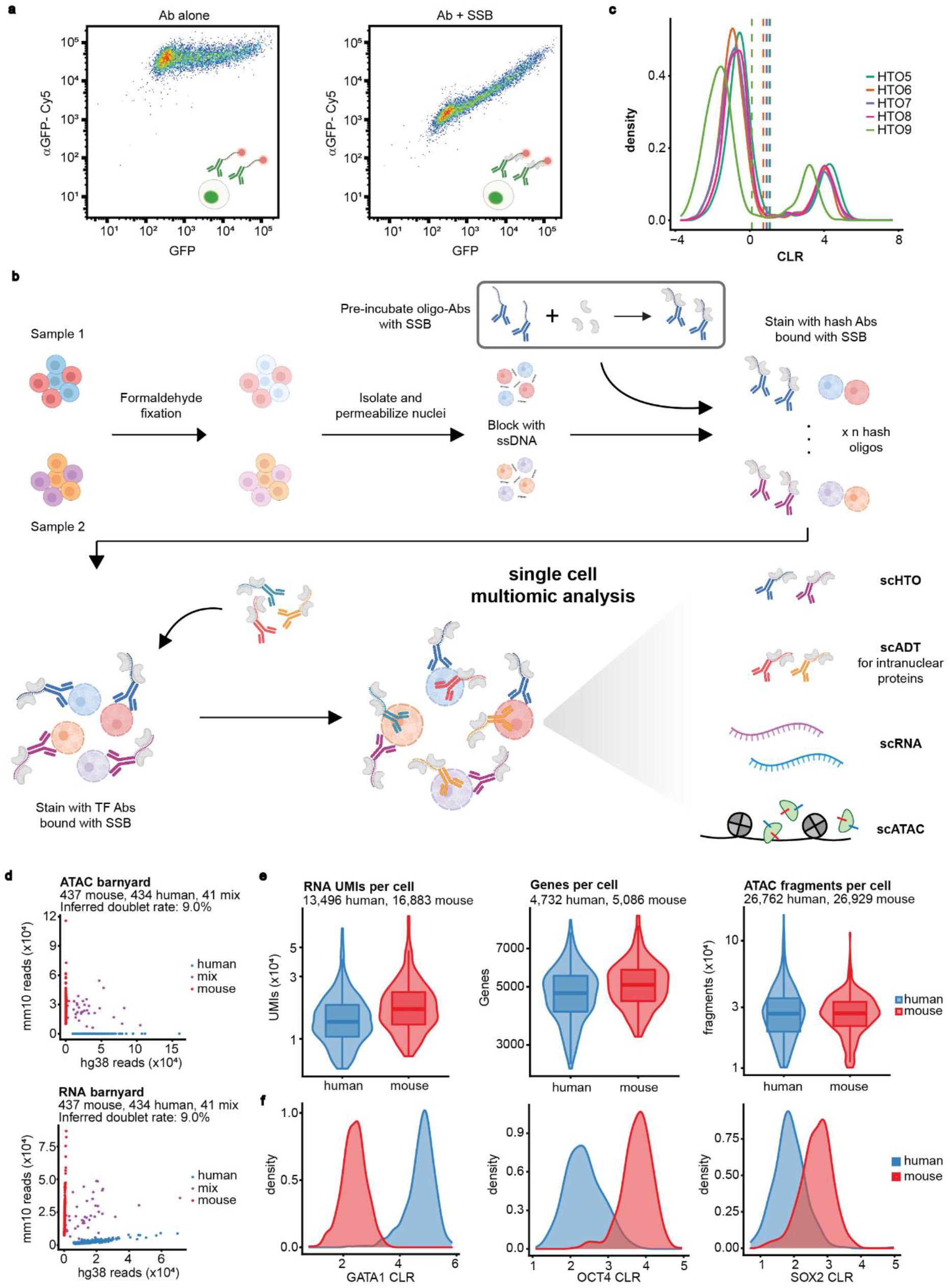
An optimized intra-nuclear staining protocol using oligo-antibodies enables simultaneous profiling of nuclear protein, chromatin accessibility and RNA transcripts in single cells. **a)** Flow cytometry plot of HEK293T cells expressing nuclear-localized GFP and stained with an anti-GFP antibody linked to an 80 bp single stranded DNA oligo with 3’-Cy5 modification. **b)** Schematic of NEAT-seq workflow. **c)** Distribution of centered log ratio (CLR) transformed counts of HTOs from anti-NPC antibodies (HTO5-9). **d)** Scatterplot of number of reads mapping to the human vs mouse genome in each cell after removing HTO doublets, with each cell colored by its classification as a human cell, mouse cell, or mixed species doublet. **e)** Distribution of RNA UMIs, genes, and ATAC-seq fragments per cell. **f)** CLR-transformed counts of ADTs corresponding to GATA1, OCT4, and SOX2 in cells classified as human or mouse cells based on mapping of ATAC-seq and RNA-seq reads to each genome.

We next sought to combine nuclear protein quantification with scATAC-seq and scRNA-seq using the 10X Genomics Multiome kit. Because cell fixation increases the doublet rate in droplet-based single cell technologies^4^, we performed a species mixing experiment with hashtag oligos^14^ (HTOs) to assess doublet identification while also evaluating the ability of our protocol to accurately measure nuclear protein abundance. We assayed a 1:1 mixture of 1,013 human K562 cells and mouse ESCs stained for mutually exclusive TFs (GATA1 for K562, and SOX2 and OCT4 for ESCs) and a nuclear pore complex (NPC) antibody linked to distinct HTOs (Fig. 1b). Clear separation between positive and negative staining for HTOs allowed us to set stringent cutoffs for positive staining and remove droplets that were positive for more than one hashing oligo and thus likely contained a doublet (Fig. 1c). Though the doublet rate was higher than for unfixed cells, the use of HTOs allowed for in silico removal of almost 50% of these doublets (Fig. 1d, Supplementary Fig. 1d, e). This hashing procedure also allows for straightforward sample multiplexing, enabling independent samples to be labeled with distinct hashing oligos, then mixed into the same experiment for hash-based downstream deconvolution.

For human and mouse singlets, we detected a median of 13,496 and 16,883 RNA UMIs; 4,732 and 5,086 genes; and 26,762 and 26,929 ATAC fragments per cell, respectively (Fig. 1e). The fragment length distribution and average TSS enrichment were also comparable to bulk ATAC-seq libraries (Supplementary Fig. 1f, g). Importantly, we observed considerable enrichment of GATA1 and OCT4 antibody-derived tag (ADT) counts and moderate enrichment of SOX2 ADT counts in their respective cell types (Fig. 1f). Together, these results show that we can simultaneously quantify endogenous nuclear protein abundance, chromatin accessibility, and gene expression in fixed single cells. We refer to this method as NEAT-seq (Nuclear protein Epitope, chromatin Accessibility, and Transcriptome sequencing).

We next applied NEAT-seq to profile primary human CD4 memory T cells. This population of cells comprises distinct memory helper T cell subsets driven by known master TFs, providing a diverse system for dissecting the regulatory mechanisms upstream and downstream of these TFs to control cell state. Our antibody panel targeted TFs that drive Th1 (Tbet), Th2 (GATA3), Th17 (RORγT), and Treg (FOXP3 and Helios) cell fate^19^. After filtering for high quality cells and removing HTO doublets, we identified contaminating CD8 memory T cells in our sample by projecting our cells onto a UMAP of peripheral blood mononuclear cells^20^ and removed these from analysis. The resulting 8,472 cells had a median TSS enrichment of 19.0 and we detected a median of 4,704 ATAC-seq fragments, 1,144 genes, and 1,999 RNA UMIs per cell.

We identified seven clusters in the population using either scATAC-seq or scRNA-seq data (Fig. 2a, Supplementary fig. 2a). Of these, we identified four clusters that contained strong enrichment for TF protein measurements, suggesting correspondence to Th1, Th2, and Th17 memory cells as well as Tregs (Fig. 2c, Supplementary fig. 2c). In most cases, these clusters also had the highest RNA levels for the respective ADT-targeted TFs, and the highest chromatin accessibility around these TF gene loci (Fig. 2c, Supplementary Fig. 2b). These clusters also showed greater accessibility at the gene loci of cytokines produced by the corresponding T cell subsets, while expressing low or undetectable levels of RNA for these cytokines (Supplementary Fig. 2g, h). This observed epigenetic priming of cytokine genes is consistent with the primed status of memory T cells^21^.

**Fig. 2:**
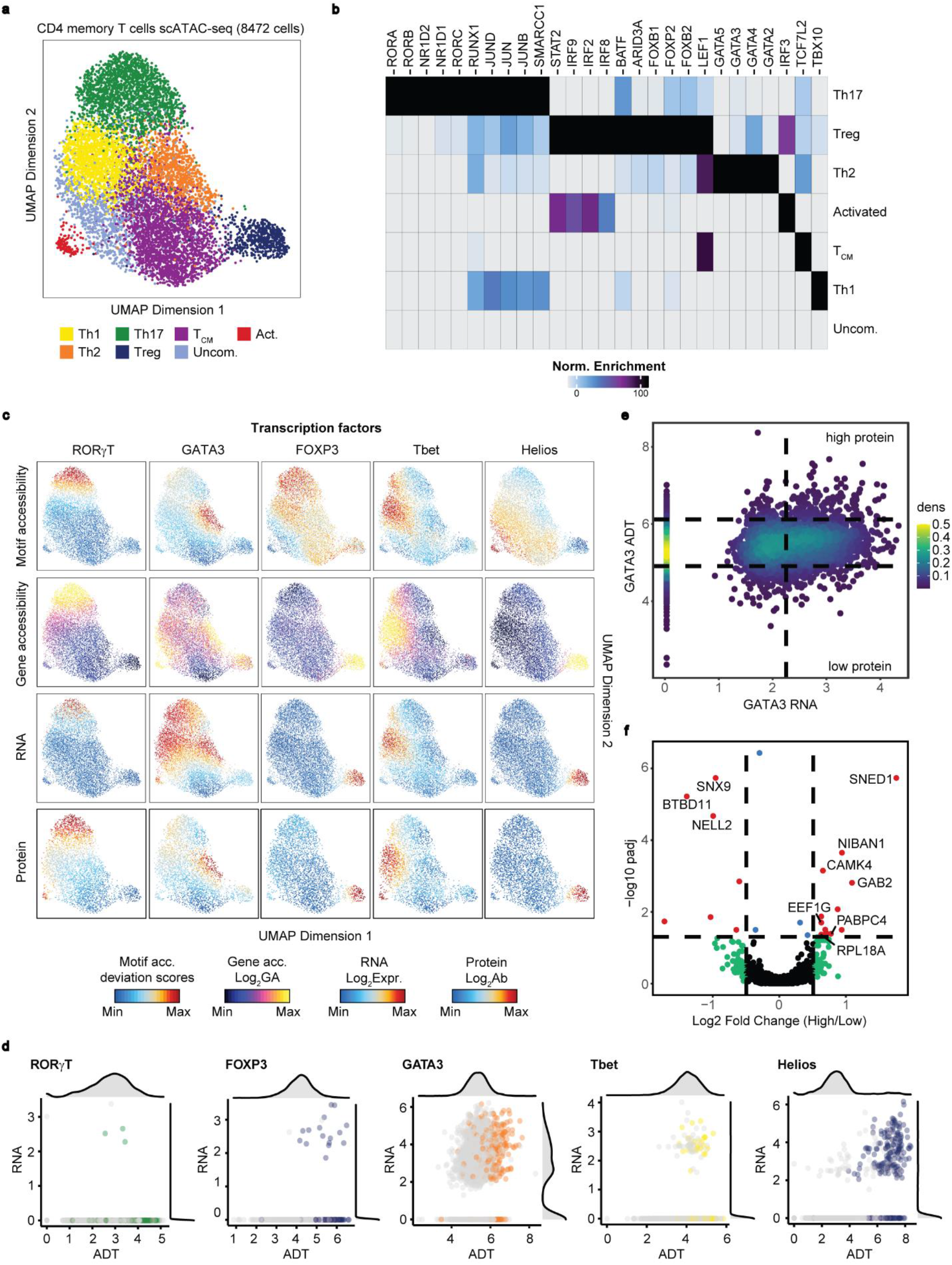
Profiling of CD4 memory T cells using NEAT-seq reveals translational regulation of GATA3. **a)** scATAC-seq UMAP of CD4 memory T cells with cell type classifications. T_CM_ = central memory, Act. = recently activated cells, Uncom. = uncommitted memory cells. **b)** Top enriched motifs in peaks that are more accessible in each cluster. **c)** Plots on scATAC UMAP of TF chromVAR deviations (motif accessibility), accessibility surrounding the TF gene locus (gene accessibility), RNA levels, and protein levels as measured by ADTs for the indicated TFs. **d)** Scatterplots with marginal histograms of log_2_-transformed read-normalized RNA vs log_2_-transformed NPC-normalized ADT counts for each TF. Colored data points represent cells belonging to the scATAC-seq cluster most enriched in expression of the indicated TF. **e)** Scatterplot of log_2_-transformed, normalized RNA vs ADT counts for GATA3 with cutoffs shown for high RNA, high protein, and low protein indicated. **f)** Differentially expressed genes between cells with high RNA and high protein vs high RNA and low protein for GATA3. Points in red represent genes with adjusted p-value < 0.05 and log_2_ fold change > 0.5.

In addition to these four T cell subtypes we identified using ADT TF measurements, we identified a cluster with increased motif accessibility for the naïve and central memory (CM) TFs, Lef1 and Tcf7^22^ (Fig. 2b, Supplementary Fig. 2d), and higher expression of the CM surface marker, CCR7^23^ (Supplementary Fig. 2e). We annotated this cluster as CM cells, although the surrounding Th1, Th2, and Th17 clusters likely also include both CM and effector memory (EM) cells, forming a continuous “effectorness gradient” that branches out from the CM cluster into EM cells of each helper T cell subtype^24^. We also identified a small cluster of cells expressing higher levels of various activated T cell markers, including CCL5, a marker of late activation^25^, and IFIT2^26^, IFIH1^27^, and OASL^28^ – genes induced during an immune response (Supplementary Fig. 2f). We therefore concluded that these cells were likely recently activated cells. Lastly, we observed a cluster lacking significant motif enrichment for any TFs. We hypothesized that these cells could represent uncommitted or virtual memory cells, a previously described memory cell type that arises without being stimulated by foreign antigen^29,30^ However, it remains possible that these cells could belong to other known T cell subsets that are unidentified here, such as Th9 or follicular helper T cells.

The use of ADTs greatly improved detection of the target TFs in a cell compared to RNA data. While smoothing the RNA signal across cells was sufficient for identification of cell types (Fig. 2c), when we plotted unsmoothed RNA-seq and ADT counts we found that only the unsmoothed ADT data were sufficient to clearly label the appropriate cell cluster while, with the exception of Helios, the RNA counts were too sparse to do so (Supplementary Fig. 2j). We observed zero RNA transcripts for the TF in over 90% of cells in the corresponding cell type for RORγT, Tbet, and Foxp3, and no transcript in over 20% of cells for GATA3 and Helios (Fig. 2d). By contrast, we observed under 0.5% dropout for the corresponding ADT counts and appropriate ADT signals were strongly enriched in the relevant cell types (Fig. 2d, Supplementary Fig. 2c).

By integrating these quantitative measurements of TF abundance with our ATAC-seq or RNA-seq data, we can identify distinct modes of TF regulatory activity. One long-standing hypothesis in the field has been that TF binding at target enhancers can lead to nucleosome eviction and creation of accessible chromatin^9^. Computational methods such as transcription factor footprint analysis and ChromVAR seek to infer this TF activity by examining global changes in accessibility patterns at TF binding sites^31,32^, but it has not been shown whether these inferences actually correspond to TF protein abundance at a single cell level. NEAT-seq revealed clear correspondence between TF abundance and ChromVAR motif accessibility scores for RORγT, GATA3, and Tbet across cells (Fig. 2c). In contrast, Helios and FOXP3 motif accessibility did not correlate with protein expression. In the case of Helios, we believe that this is due to a “collision” of binding motifs: “GGAA” is the core motif for Helios, which is highly similar to the NFAT motif, a previously described binding partner for Helios^33^. We hypothesized that this Helios motif denotes Helios binding driven by interactions with NFAT in GM12878 where the two TFs are co-expressed and from which the ChIP-seq motif is derived, and is therefore not representative of the binding motif in Tregs. Supporting this hypothesis, we found that NFAT expression and NFAT motif accessibility were highly overlapping with accessibility of the B cell-derived Helios motif in CD4 memory cells, and that NFAT expression was low in Tregs (Supplementary Fig. 2i). In contrast, the lack of concordance between FOXP3 expression and motif accessibility is consistent with previous studies showing that FOXP3 binds to pre-existing enhancers to drive Treg fate^34^, indicating that FOXP3 binding relies on the chromatin remodeling activity of other TFs. These observations suggest that, while three of the TFs in our panel appear to promote accessibility at their binding sites, FOXP3, and possibly Helios, appear unable to directly influence the chromatin accessibility landscape at their putative binding sites.

TF protein abundance measurements also enable us to identify post-transcriptional regulatory processes that would not be observable through scRNA-seq alone. With GATA3, we detected clear discordance between protein levels and RNA expression across cells, with high GATA3 RNA expression observed across several memory T cell subsets and high GATA3 ADT levels observed only in the Th2 cluster (Fig. 2c). These ADT differences, but not RNA differences, were correlated with global changes in GATA3 motif accessibility, suggesting that ADTs faithfully report on chromatin modulating potential of this TF. Conversely, accessibility proximal to the GATA3 gene was increased in a range of cells in a pattern similar to GATA3 RNA expression. These observations are consistent with post-transcriptional regulatory mechanisms restricting GATA3 protein expression in memory T cells.

To identify genes that may be involved in post-transcriptional GATA3 regulation, we first identified cells expressing high levels of GATA3 RNA, then performed differential expression analysis between cells with high vs low levels of GATA3 ADT using Seurat (Fig. 2e). Among the top upregulated genes (FDR < 0.05) were several core translation regulators, including the elongation factor EEF1G, large ribosome subunit RPL18, and poly-A binding protein PABPC4, as well as more indirect regulators such as NIBAN1, which promotes translation by regulating phosphorylation of the initiation factors EIF2A and EIF4EBP1^35^ (Fig. 2f). GATA3 translation is regulated by PI3K signaling through mTOR^36^ which, like NIBAN1, phosphorylates EIF4EBP1 to allow assembly of the initiation complex^37^. We also observed upregulation of a direct activator of PI3K, GAB2, in cells with high GATA3 protein levels. These results suggest that upregulation of genes that promote translation may play a role in driving GATA3 protein production in the Th2 subset of memory T cells. Together, our results show that the TFs in our panel can be separated into three categories based on the primary mechanism used to modulate their activity: those regulated at the level of transcription and show concordant RNA, protein, and motif accessibility patterns (RORγT, and Tbet), those regulated transcriptionally but that require other TFs to bind chromatin (Helios and FOXP3), and those regulated post-transcriptionally (GATA3).

Simultaneous measurement of nuclear TF protein abundance, chromatin accessibility, and gene expression also enables the correlation of each of these data types to uncover regulatory links between trans-acting factors, the cis-acting elements they bind, and resulting changes in expression. Such an analysis is challenging with only RNA-seq data, as low TF transcript counts and post-transcriptional regulation of TF abundance make correlation both technically and biologically problematic. We found hundreds of cis-regulatory elements with accessibility significantly correlated with the protein levels of RORγT, Tbet, and GATA3 (FDR < 0.05). As expected, the corresponding TF motif was significantly enriched in these peaks (Supplementary Fig. 3a). We observed no significant enrichment for the FOXP3 and Helios motifs in correlated peaks, consistent with our earlier observations that these TFs are not correlated with global accessibility changes (Fig. 2c).

We similarly searched for genes associated with each TF and identified dozens of genes with RNA expression significantly correlated with protein levels of each TF (Supplementary Fig. 3b). Within these correlated gene sets were genes known to be enriched or functionally important in the memory T cell subset driven by the TF in question, such as IL4R for GATA3 and CTLA4 for both Helios and FOXP3^38^. We also explored how differences in TF protein abundance within a cluster might reveal heterogeneity in cell state and detected a modest difference in RORγT motif accessibility between cells containing high and low RORγT protein levels within the annotated Th17 cluster (Supplementary Fig. 3d). RORγT protein levels within this cluster were significantly correlated with a number of genes known to function in Th17 cells (RORA^39^, PTPN13^40^, ZBTB16^40,41^; Supplementary Fig. 3e), while genes anti-correlated with RORγT levels included genes enriched in naïve CD4 cells (NELL2, MAML2)^42^ and genes that promote cell cycle (YPEL5, CCND3)^43,44^, suggesting that cells with low RORγT in the Th17 cluster may represent a more proliferative precursor in memory cell differentiation.

To identify candidate genes directly regulated by each TF through a TF-associated enhancer, we overlapped the top TF ADT-correlated genes with top TF ADT-correlated peaks containing the corresponding TF motif that were within 100 kb of the gene promoter and filtered for significant peak-gene linkages (Fig. 3a). We performed this analysis for the TFs that showed correlation between TF abundance and motif accessibility and identified 167 candidate TF-peak-gene linkages for GATA3, 345 for RORγT, and 81 for Tbet (Fig. 3b). Included in these candidate TF targets were canonical surface markers for the corresponding cell type: Among the GATA3 targets were Th2 markers CCR4, CCR8, and IL4R, and among RORγT targets was the Th17 marker, CCR6 (Fig. 3c, Supplementary Fig. 3c).

**Fig. 3:**
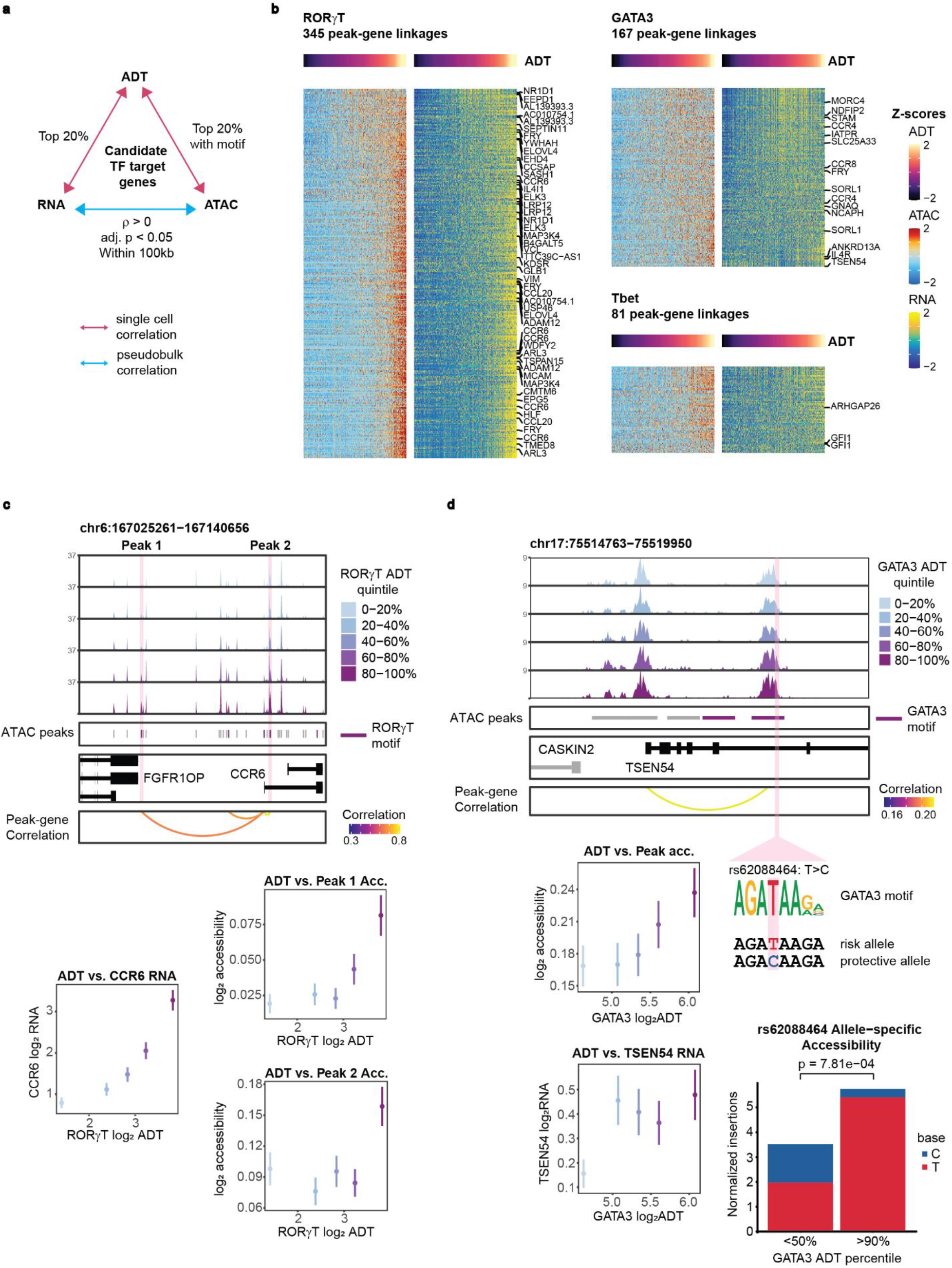
Identification of peak-gene linkages associated with master TF protein expression. **a)** Diagram showing criteria used to identify candidate TF-peak-gene linkages for each TF. **b)** Heatmaps of peak-gene linkages correlated with abundance of the indicated TF identified based on criteria in (a). Genes that are in the top 200 genes significantly enriched in the T cell subset driven by the corresponding TF based on data by Schmiedel et al.^38^ are labeled. **c)** Top: CCR6 ATAC-seq tracks in CD4 memory cells separated into quintiles by RORγT ADT levels, along with significant peak-gene linkages (adj. p < 0.05). Peaks containing a RORγT motif are indicated. Bottom: CCR6 RNA expression and accessibility at the highlighted RORγT motif-containing peaks as a function of RORγT ADT levels. Mean is shown with standard error of the mean. **d)** Top: TSEN54 ATAC-seq tracks as in (c), but for GATA3 ADT, with the indicated GWAS SNP location highlighted. Bottom left: TSEN54 RNA expression and accessibility at the highlighted SNP-containing peak as a function of GATA3 ADT levels. Mean is shown with standard error of the mean. Bottom right: Read-normalized Tn5 insertions per cell per million reads mapping to the risk vs protective allele in cells above the 90th percentile vs cells below the 50th percentile of GATA3 ADT levels. P-value is calculated using a one-sided binomial test on T:C ratio of insertion counts.

We also reasoned that the TF-peak-gene linkages we identified could be used to interpret the effects of non-coding GWAS SNPs on TF activity and connect them to putative target genes. We overlapped peaks in our TF-peak-gene linkages with candidate causal GWAS SNPs^45^ and identified a SNP, rs62088464, located within a GATA motif sequence in a GATA3 ADT-associated peak. The risk allele, which preserves the GATA motif, is associated with decreased pulmonary function as measured by decreased forced vital capacity^46^, which can result from pulmonary fibrosis and several other inflammatory lung diseases associated with Th2 immune responses^47^. The gene linked to the peak containing this SNP encodes the tRNA splicing endonuclease, TSEN54, a gene with significantly enriched expression in the sputum of patients with type-2 airway inflammation^48,49^ (Fig. 3d). Since our T cell donor was heterozygous for this SNP, we examined whether the risk allele was more accessible than the protective allele in cells with high GATA3 protein levels. Indeed, we observed that almost all ATAC-seq reads in the top 10% of cells ranked by GATA3 ADT levels mapped to the risk allele, while this difference was far less pronounced in cells with lower levels of GATA3 ADT (Fig. 3d, p = 7.81×10^−4^). Together, these results suggest that GATA3 binds the risk allele sequence to activate the regulatory element and drive expression of TSEN54 and that this binding is disrupted with the protective allele.

NEAT-seq provides a new avenue for studying the quantitative effects of epigenetic regulator abundance on both chromatin and gene expression state in primary human samples. Whereas previous studies investigating dosage-dependent effects of TFs often required building cell lines with a combination of hypomorphic and null alleles^50,51^ or inducible expression systems^52^, we demonstrate that NEAT-seq can measure the molecular consequences of continuous changes in TF levels in a biologically relevant setting for a panel of proteins simultaneously. Measuring nuclear protein levels using ADTs is more sensitive and accurate than RNA measurements for both technical and biological reasons. Since nuclear proteins encompass many proteins involved in gene regulation including TFs and chromatin modifiers, the capacity to link nuclear protein levels to epigenetic and transcriptional status provides a powerful approach for studying gene regulation. While oligo-antibodies against nuclear proteins are currently limited, we anticipate that these will become more readily available as demand increases. Incorporating additional modalities such as cytoplasmic and cell surface proteins, CRISPR gRNA sequencing, and TCR sequencing will enable measurement of the effects of cellular perturbations and signaling pathways on cell state, providing an even more comprehensive picture of cellular programs. By measuring how these modalities change in relation to each other in dynamic processes such as cellular differentiation, reprogramming, transformation, and activation, we will be able to dissect the molecular regulatory mechanisms that drive cell fate determination and disease states in a variety of systems.

## Acknowledgements

We thank Reema Baskar, Sarah Pierce, Sandy Klemm, YeEun Kim, and members of the Greenleaf lab for helpful discussions and suggestions. We also thank the Stanford FACS facility and Stanford Functional Genomics Facility for technical support. Figure schematics were created with BioRender.com. This work was supported by funding from the NIH (P50HG007735, R01HG008140, R01HG00990901, U19AI057266, and UM1HG009442), the Rita Allen Foundation, the Baxter Foundation Faculty Scholar Grant and the Human Frontiers Science Program (RGY006S) to WJG. WJG is a Chan Zuckerberg Biohub investigator (grant # 2017-174468 and 2018-182817). Fellowship support was provided by the Stanford School of Medicine Dean’s Fellowship and NIH (F32GM135996) to AFC and a training grant from the National Institute of Standards and Technology to BP.

## Author Contributions

AFC and WJG conceived the project and designed the experiments with input from all authors. AFC led method development and performed experiments with help from ASK. AFC and BP performed bioinformatic analysis, visualization and interpretation. AFC and WJG drafted the manuscript, and AFC, BP, BOR, and WJG revised and edited the manuscript with input from all authors.

## Experimental methods

### Cell culture

Mouse V6.5 ESCs were cultured on gelatin-coated plates in Knockout DMEM (Thermo Fisher #10829018) supplemented with 7.5% ES-qualified serum (Applied Stem Cell #ASM-5017), 7.5% Knockout Serum replacement (Thermo Fisher #10828-028), 2mM L-glutamine (Gibco #35050061), 10mM HEPES (Gibco #15630080), 100 units/mL penicillin/streptomycin (Gibco #151401222), 0.1mM non-essential amino acids (Gibco #11140050), 0.1mM beta-mercaptoethanol (Gibco #21985023), and LIF. The human chronic myeloid leukemia cell line, K562, was purchased from ATCC and cultured in RPMI 1640 medium (Gibco #11875119) containing 15% FBS and 100 units/mL penicillin/streptomycin. HEK 293T cells were cultured in DMEM with GlutaMAX (Gibco #10566024) containing 10% FBS and 100 units/mL penicillin/streptomycin. Frozen vials of primary human CD4+CD45RO+ memory T cells were purchased from STEMCELL Technologies (Cat #70031).

### GFP transfection, staining, and sorting

On day 0, HEK 293T cells were seeded at 4 million cells per 10cm plate. On day 1, cells were transfected with 6ug nuclear-localized GFP construct (Addgene #67652) using Fugene HD transfection reagent (Promega). Cells were harvested and stained using anti-GFP antibody (Biolegend 338002) linked to an 80 bp ssDNA oligo with 3’ Cy5 fluorophore as described in the oligo-antibody staining methods section, except without RNase inhibitor or DTT. A control stain was performed with the oligo-antibody in the absence of SSB. Stained cells were resuspended in PBS and analyzed on an LSRII flow cytometer or sorted on a BD FACS Aria II.

### Antibody conjugation

Antibodies were conjugated with streptavidin using the Lightning-Link Streptavidin Conjugation Kit from Abcam (ab102921) according to manufacturer’s instructions. NaCl and Tween were added to the conjugated antibody mixture to a final concentration of 0.5M NaCl and 0.01% Tween and mixed with biotinylated oligos (purchased from IDT) at equimolar ratio. The mixture was incubated overnight at room temperature and unbound oligo was removed using Amicon 100KDa centrifugal filters (UFC510008). Antibody conjugates were eluted and stored in PBS. Antibodies conjugated with streptavidin were GATA1 (Abcam ab241393), OCT4 (R&D AF1759), SOX2 (R&D MAB2018), nuclear pore complex antibody (Biolegend 902901), and GFP (Biolegend 338002). Antibodies in the TF panel for CD4 memory T cells were directly conjugated to oligos by BD Biosciences. The antibodies in the panel were the following clones from BD Biosciences: GATA3 (L50-823), Tbet (4B10), RORγT (Q21-559), FOXP3 (259D/C7), and Helios (22F6).

### Binding of single stranded DNA binding protein to oligo-antibodies

To bind EcoSSB (Promega M3011) to the antibody-oligos, we incubated the antibody and EcoSSB in 50ul of 1X NEBuffer 4 for 30 min at 37 degrees Celsius. We then added a final concentration of 3% BSA, 1X PBS, and 1U/ul RNase inhibitor to the antibody mix in a final volume of 100ul for staining cells. To calculate the amount of EcoSSB needed to saturate binding sites on the antibody oligos, we estimated that each antibody was conjugated to an average of 2 oligos of 95bp, and each EcoSSB tetramer would bind with a ~35bp footprint^53,54^, requiring 6 EcoSSB tetramers per antibody. Based on the concentration of antibody being used and reported K_d_ of EcoSSB (in the ~2nM range)^17^, we can then estimate the amount of EcoSSB necessary to bind a given fraction of oligos (aiming for > 0.9) using the following equation:

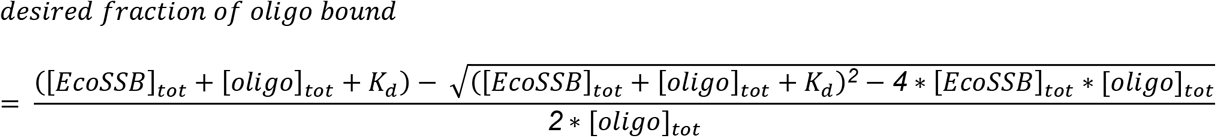

where [oligo]_tot_ = antibody concentration * 2 oligos * 3 EcoSSB binding sites per oligo.

### Oligo-antibody staining

Cells were fixed in 1.6% formaldehyde in PBS for 2 min at room temperature, then quenched with 0.25M glycine for 5 min on ice and spun down at 600g for 5 min. Cells were washed twice with PBS and then resuspended in lysis/permeabilization buffer (20mM Tris-HCl pH 7.5, 150mM NaCl, 3mM MgCl2, 0.5% NP40, 0.1% Tween-20, 0.01% digitonin, 1U/ul RNase inhibitor, 1mM DTT). Cells were incubated on ice for 10 mins, pelleted at 600g for 5 mins, and washed twice with wash buffer (20mM Tris-HCl pH 7.5, 150mM NaCl, 3mM MgCl2, 0.1% Tween-20, 1U/ul RNase inhibitor, 1mM DTT). Cells were incubated in staining buffer (PBS with 3% BSA, 1U/ul RNase inhibitor) with 1mM DTT and 1mg/ml of single stranded DNA (ssDNA) for 30 mins at room temperature, pipetting often to resuspend cells. For the flow cytometry experiments involving GFP staining, salmon sperm DNA was used for the ssDNA block. However, due to significant amounts of annealing to form double stranded DNA that would result in contaminating reads in ATAC-seq data, we switched to using either a mixture of random 30-mers or a 30bp ssDNA oligo sequence with no complementarity to the mouse or human genome for multiome experiments. To ensure no priming would occur with these oligos, they were modified with a terminal dideoxy cytosine.

After blocking with ssDNA, Tween was added to a final concentration of 0.1% and cells were pelleted and washed once with staining buffer + 0.1% Tween. Cells were then split into 5 tubes and each tube of cells was incubated with an anti-NPC antibody linked to a distinct HTO (pre-bound with SSB) for 30 min at room temperature. Cells were washed twice with staining buffer + 0.1% Tween, re-pooled, and incubated with TF antibody mix for 30 min at room temperature. For the CD4 memory T cell experiment, cells were split into two tubes prior to incubating with two concentrations of the TF antibody mix. A distinct hashing antibody was also added to the two TF antibody mixes to identify the concentration of antibody that each cell was stained with. Cells were then washed twice with staining buffer + 0.1% Tween, and cells incubated with different concentrations of TF antibody were pooled. Cells were washed once more with PBS containing 1% BSA and 1U/ul RNase inhibitor, then resuspended in 1X Nuclei buffer containing 1U/ul RNase inhibitor from the 10X Genomics Multiome kit. The cell suspension was then filtered through a 40um Flowmi strainer 2-3 times until nuclei clusters were removed.

### Antibody concentrations

The NPC, GATA1, SOX2, and OCT4 antibodies were all used at 0.3ug in 100ul of staining buffer (3ug/mL). The two antibody concentrations for TF antibodies used in the CD4 memory T cell experiment are indicated below:

| **Antibody** | **Conc. 1 (ug/mL; marked by NPC1)** | **Conc. 2 (ug/mL; marked by NPC2)** |
| --- | --- | --- |
| RORγT | 0.39075 | 1.95375 |
| Foxp3 | 2.5 | 5 |
| GATA3 | 3.125 | 15.625 |
| Helios | 0.39075 | 1.95375 |
| Tbet | 3.125 | 15.625 |

### Single cell library preparation and sequencing

Antibody-stained cells in 1X Nuclei buffer were processed using the 10X Genomics Multiome kit as indicated in the standard protocol (Rev A) to generate ATAC-seq and RNA-seq libraries. For the barnyard experiment, 1,500 cells were targeted in one lane of the chip. For the CD4 memory T cell experiment, 6,000 cells were targeted per lane and 2 lanes were used. During the pre-amplification step, Truseq read 2 (CAGACGTGTGCTCTTCCGATC) and Nextera read 2 (GGCTCGGAGATGTGTATAAGAGACAG) primers were spiked in at 0.2uM final concentration to amplify ADT and HTO oligos. To generate ADT and HTO libraries, 35ul of pre-amplification product from step 4.3p was amplified with indexing primers using 2X NEB Next High-Fidelity PCR Master Mix (M0541). A double-sided SPRI bead clean up was performed using 0.6X SPRI beads (retaining supernatant) and then adding additional SPRI beads to a final concentration of 1.2X, washing with 80% ethanol, and eluting ADT or HTO libraries from beads using EB buffer. Libraries were quantified by PCR using a PhiX control v3 (Illumina FC-110–3001) standard curve. scATAC-seq libraries were sequenced alone on a NextSeq 550 sequencer and ADT libraries were either sequenced alone on a MiSeq (for the barnyard experiment) or together with scRNA-seq libraries on a NextSeq 550 (for the CD4 memory T cell experiment). Recommended sequencing read configurations for 10X Multiome libraries were used for scATAC- and scRNA-seq libraries. For sequencing of the ADT libraries from the barnyard experiment, the read configuration was 28bp Read 1, 48bp Read 2, and 8bp for Index 1 and 2. We sequenced approximately 300,000 read pairs per cell for both scATAC-seq and scRNA-seq libraries and 7,000 read pairs per cell for ADT libraries in the barnyard experiment. We sequenced approximately 40,000 read pairs per cell for scATAC-seq, 35,000 read pairs per cell for scRNA-seq libraries, and 5,000 read pairs per cell for both the ADT and HTO libraries in the CD4 memory T cell experiment.

### Antibody oligo sequences

ADT oligos and HTO oligos from the barnyard experiment had a partial Truseq read 2 sequence followed by 12bp UMI, 36bp antibody-specific barcode, and 25bp poly A tail as follows:

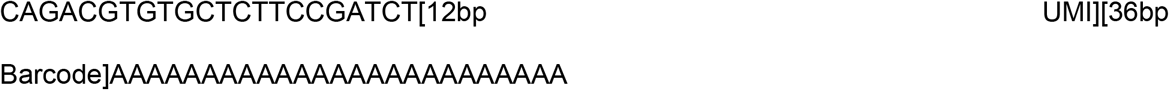

HTOs for the CD4 memory T cell experiment were similarly designed, except they instead had a partial Nextera read 2 sequence to allow separate amplification of TF antibody oligos from HTOs, which often stain at higher levels:

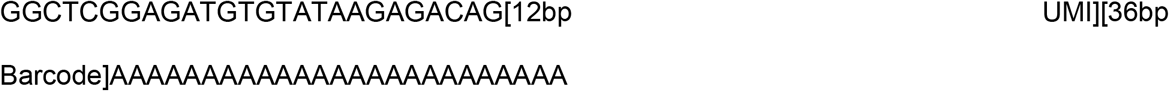

Note that the hashing antibody used together with the TF antibody panel for marking the two antibody concentrations tested in CD4 memory T cells was linked to an ADT oligo with a partial Truseq read 2 sequence so that it would be amplified with the TF ADTs and could be used to normalize TF ADT counts. All antibody barcode sequences are provided in Supplementary Table 1.

## Analytical methods

### ADT and HTO processing

Raw sequencing data were converted to fastq format using bcl2fastq (Illumina). ADTs and HTOs were then assigned to individual cells and antibodies using the matcha barcode matching tool^55^. Cell barcodes were matched based on exact matches, and up to 3 mismatches were allowed in antibody barcode sequences. Counts for each antibody were tabulated by counting UMIs. In the barnyard experiment, cells with fewer than 200 ADT+HTO UMIs were excluded from analysis. In the CD4 T cell experiment, cells with fewer than 75 HTO UMIs or 100 ADT UMIs were excluded. All HTO count data and TF ADT count data from the barnyard experiment were normalized using a centered log ratio (CLR) transformation as previously described^1^. For the CD4 memory T cell experiment, TF ADT counts were normalized to HTO counts from the anti-NPC HTO that was added to distinguish two different concentrations of the TF antibody panel used to stain cells, since we expected that levels of the nuclear pore complex should be relatively constant across cells. We then multiplied by 250 (i.e roughly the median number of NPC counts per cell), added one pseudocount, and log_2_-transformed counts. We chose the NPC normalization method because it was more robust than CLR transformation in cases where cells are primarily positive for only one antibody in the panel, as was the case for the CD4 memory T cells.

### Doublet detection using HTOs

For doublet detection in the barnyard experiment, we filtered for cells with at least 400 total ADT counts, and performed CLR-transformation on HTO counts only. CLR cutoffs for positive staining of each HTO was performed automatically in a similar manner to Seurat’s HTODemux function^56^. Cells were k-means clustered based on CLR-normalized HTO counts, with the k equal to the number of hashing oligos. This serves as a rough HTO assignment which can be used to infer background staining distributions. The cutoff value for each HTO was determined by taking the 99^th^ percentile of a normal distribution fit to the CLR-normalized HTO counts in the k-1 clusters with the lowest mean value for the given HTO. This differs from Seurat’s HTODemux by using a normal distribution on CLR-normalized counts rather than a negative binomial distribution on raw counts, and in using the bottom k-1 clusters to fit the background distribution rather than the bottom 1 cluster. After computing cutoffs for each HTO, we removed cells which were not positive for exactly 1 HTO, annotating the cells positive for >1 HTO as doublets.

For doublet detection in the CD4 memory T cell experiment, we filtered for cells with at least 75 HTO counts per cell and performed CLR-transformation on HTO counts only. We set CLR cutoffs for positive staining of each HTO individually based on the bimodal distribution for each HTO and only cells positive for exactly one HTO were retained. Since we also incorporated two hashing oligos in the TF staining step to distinguish between two antibody concentrations used, we also annotated doublets using these HTOs and removed them from analysis.

### Barnyard experiment species analysis

Raw sequencing data were converted to fastq format, and aligned to a chimeric hg38 and mm10 reference genome using cellranger-ARC v1.0.1 from 10x Genomics. First, we filtered droplets for high-quality cells based on >7,500 RNA UMIs, >10,000 unique ATAC fragments, and TSS enrichment > 10. TSS enrichment was calculated using the combined set of mouse + human TSS coordinates and the default parameters of ArchR’s TSS enrichment. Next, we annotated species based on the fraction of reads aligning to either the mouse genome or the human genome. For ATAC-seq reads, this cutoff was manually set to >95% of reads aligning to a single species. For RNA-seq reads we observed greater cross-cell read contamination, particularly from mouse transcripts which had high abundance in non-cell droplets. As a result, we set a cutoff of >70% reads aligning to the human genome, or >95% reads aligning to the mouse genome. For our main doublet analysis, we considered cells to be mouse-human doublets if they did not pass the species cutoff for both their ATAC-seq and RNA-seq reads.

Inferred doublet rates were calculated by dividing the observed doublet rate by the fraction of cell pairings expected to be between mouse and human cells 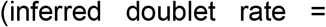 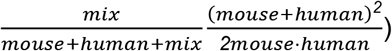. For perfectly even mixtures of mouse and human cells, the inferred doublet rate will be twice the observed doublet rate, and deviations from even mixtures will increase the inferred doublet rate relative to the observed doublet rate.

### scATAC-seq analysis

Raw sequencing data were converted to fastq format and aligned to the hg38 reference genome using cellranger-ARC v.1.0.1 from 10X Genomics. Fragment files were then loaded into ArchR (v1.0.2) using the createArrowFiles function. Cells with a TSS enrichment < 10 or fewer than 1000 unique fragments per cell were removed from analysis along with HTO-annotated doublets. Remaining cells were projected onto a reference dataset of hematopoietic cells^20^, using a liftover of the published hg19 peak coordinates to hg38 and the published LSI loadings for each peak. Cell type annotations were transferred as the most common cell type from the 10 nearest neighbors, and contaminating CD8 memory T cells were removed from further analysis. We next computed an iterative LSI dimensionality reduction using the addIterativeLSI function with the default tile matrix (insertion counts in 500bp bins across the genome) and 4 iterations. Clustering was then performed using the addClusters function and a UMAP was generated using addUMAP, both with default parameters.

To call peaks, we first generated insertion coverage files from pseudobulk replicates grouped by cluster using addGroupCoverages and then called peaks with macs2 using addReproduciblePeakSet with default parameters. We then generated a matrix of insertion counts for each peak across all cells using addPeakMatrix. To aid in cluster identification, we identified marker peaks unique to each cluster and identified TF motifs enriched in these peaks using getMarkerFeatures (useMatrix = “PeakMatrix”) and peakAnnoEnrichment. Results were plotted using plotEnrichHeatmap(enrichMotifs, n = 5, transpose = TRUE, cutOff = 5). We can also predict TF activity by measuring differences in TF motif accessibility across cells using chromVAR^31^. We first determined which peaks contain a motif of interest for motifs in the CISBP database^57^ using addMotifAnnotations with the option motifSet = “cisbp”. We then added a background peak set with similar GC content and number of fragments and computed motif deviations for all motifs using addBgdPeaks and addDeviationsMatrix, respectively.

To further help with cluster identification using ATAC-seq data, we can predict gene expression or epigenetic priming of a locus by calculating gene activity scores for each gene based on accessibility in the region surrounding the gene locus. These scores were calculated in ArchR during Arrow file creation with the option addGeneScoreMat = TRUE.

To compare RORγT TF footprinting in Th17 cells with different levels of RORγT ADT, we calculated imputed RORγT ADT values using the ArchR implementation of the MAGIC algorithm^58^ with the imputeMatrix function and selected cells within the Th17 cluster that contain the highest and lowest 30% of imputed RORγT ADT counts. We then generated insertion coverage files from these cell subsets using addGroupCoverages and calculated the RORγT TF footprint using getFootprints. Results were plotted using plotFootprints with normMethod = “Subtract”, although similar results were obtained using normMethod = “Divide”.

### scRNA-seq analysis

Raw sequencing data were converted to fastq format and aligned to the reference genome using cellranger-ARC v.1.0.1 from 10X Genomics. For each lane, the gene expression matrix from the filtered_feature_bc_matrix was used to create a Seurat object using Seurat v3.2.1. The two lanes of CD4 memory T cell data were then merged into one Seurat object and filtered for cells used in the scATAC-seq analysis. Data were normalized with NormalizeData (normalization.method = “LogNormalize” and scale.factor = 10000). For principal component analysis, we identified the top 2000 variable genes using FindVariableFeatures (selection.method = “vst”) and RunPCA was performed on scaled data using these variable features. We then clustered cells using FindNeighbors with dimensions 1:15 and FindClusters with resolution 0.6. The RNA UMAP was generated with RunUMAP using dimensions 1:15. FindAllMarkers was used to identify marker genes enriched in each cluster.

To identify candidate regulators of GATA3 translation, we added ADT data to our Seurat object using CreateAssayObject. We first filtered for cells expressing high GATA3 RNA (natural log-normalized counts > 2.25) and then identified cells expressing high GATA3 ADT (log2 NPC-normalized counts > 6.12) or low GATA3 ADT (log2 NPC-normalized counts < 4.9116 to match number of cells in high GATA3 ADT subset). To identify differentially expressed genes between these two subsets, we ran FindMarkers. We converted the natural log-based fold change values output from Seurat v3 to log2 fold changes and calculated adjusted p values using Benjamini-Hochberg correction.

### Data visualization

Unless otherwise indicated in the text, visualization of TF motif deviation Z-scores, gene activity scores, RNA, and ADTs on the ATAC UMAP embedding was done by plotting imputed values using ArchR’s plotEmbedding function. Ridge plots of normalized ADT counts and scatterplots with marginal histograms of normalized ADT vs RNA counts were generated using ArchR’s plotGroups (plotAs = “ridges”) and ggpubr’s ggscatterhist, respectively. Normalized ADT counts were calculated as log2(250*(TF ADT counts/NPC HTO counts)+1). Normalized RNA counts were calculated as log2(10000*(TF RNA counts/total UMI counts)+1).

### Identifying peaks and genes correlated with TF abundance

To identify peaks and genes with changes that correlate with TF ADT levels, Spearman correlation values were calculated between normalized ADT counts for each TF and either normalized Tn5 insertion counts or normalized RNA counts for all peaks and genes with >10 observed reads across single cells. Raw p-values for correlations were calculated in the same manner as R cor.test, namely using a two-sided t-test with n-2 degrees of freedom where 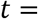 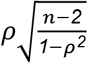 and n is the number of cells. P-values were multiple-hypothesis corrected for each ADT using the “BH” method of R’s p.adjust, and significant correlations were defined as adjusted p-value < 0.05. TF motif enrichment in significantly correlated peaks was calculated using a hypergeometric test.

### Identification of correlated peaks and genes

To identify peaks and genes where peak accessibility correlated with gene expression, we formed 500 aggregates of 100 cells each using the 99 nearest neighbors of randomly selected cells in LSI coordinates. These aggregates were constrained to have a maximum pairwise overlap of 80% of cells. Gene expression and peak accessibility for each aggregate was calculated by averaging the normalized accessibility or expression values across all cells in the aggregate. For all peak-gene pairs within 100kb of each other, we calculated Spearman correlation and significance using a two-sided t-test as for our peak-TF and gene-TF correlations.

### Identifying TF-peak-gene linkages

To identify candidate direct target genes of a TF, we identified TF ADT-correlated genes that had a TF ADT-correlated peak nearby containing the TF sequence motif. Specifically, we overlapped the top 20% of ADT-correlated genes with the top 20% of ADT-correlated peaks containing the corresponding TF motif, sorted by Spearman correlation calculated across single cells. For the overlap, we required that the peak-gene distance be less than 100kb and that accessibility of the peak and expression of the linked gene be significantly correlated (adjusted p-value < 0.05 for Spearman correlation, as described above).

### Analysis of fine-mapped GWAS variants

To identify candidate causal SNPs regulated by a TF and link the SNP to a putative target gene, we obtained a comprehensive list of fine-mapped GWAS SNPs (https://pics2.ucsf.edu/PICS2.html) and overlapped these with peaks from our identified GATA3 TF-peak-gene linkages. We focused on rs62088464, a SNP located within a GATA motif site and for which our donor was heterozygous for the risk allele. To determine allele-specific differences in accessibility at this SNP, we identified all reads overlapping this SNP with mapq > 30 using pysam’s pileup method^59,60^. To stratify cells by GATA3 expression, we z-score transformed the CLR-normalized GATA3 expression levels for each of the two antibody titration levels to ensure they were on comparable scales, then performed smoothing using the ArchR version of the MAGIC algorithm to reduce noise. Cells were divided based on their rank in the smoothed GATA3 vector. Allele-specific accessibility was determined using a one-sided binomial test, comparing the allele frequency in the top 10% of GATA3 cells using the bottom 50% as a null hypothesis.

---

**Supplementary Fig. 1:**
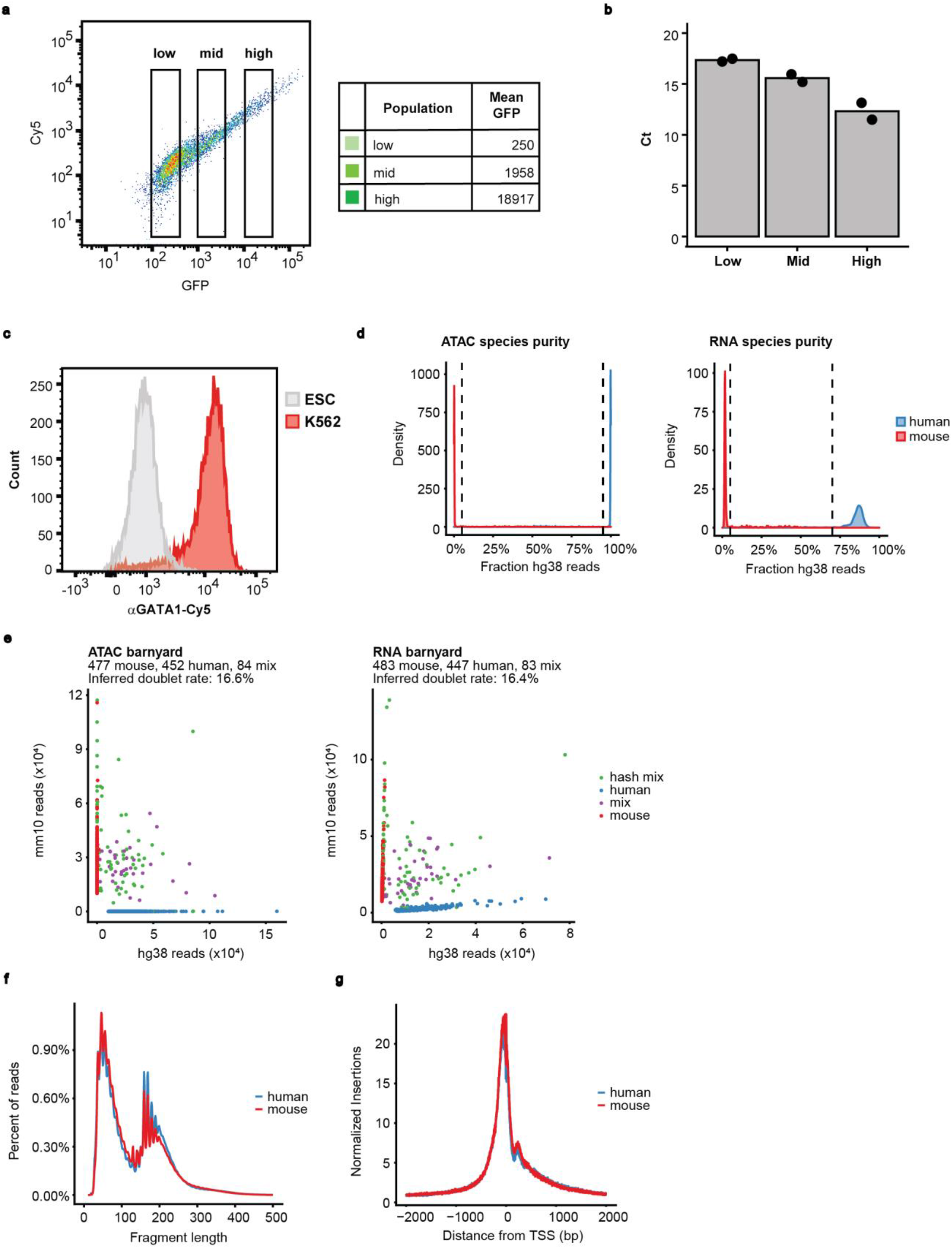
NEAT-seq enables sensitive quantification of endogenous transcription factors in single cells while generating high quality ATAC-seq and RNA-seq data. **a)** Sorting of cells expressing low, mid, or high levels of nuclear GFP that have been stained with an anti-GFP oligo-conjugated antibody. **b)** Quantitative PCR for the conjugated oligo from equal cell numbers of sorted populations in **(a)** for n = 2 technical replicates. **c)** Staining of K562 cells and mouse ESCs for endogenous GATA1 protein using an anti-GATA1 antibody linked to an 80bp oligo with 3’-Cy5. **d)** Cutoffs for annotating a cell as human, mouse, or mixed based on percentage of reads in a cell mapping to the human genome for ATAC-seq and RNA-seq data. **e)** Scatterplot of number of reads mapping to the human vs mouse genome in each cell prior to removing HTO doublets, with each cell colored by its classification as a human cell, mouse cell, mixed species doublet, or an HTO doublet. **f)** Fragment length distribution of ATAC-seq data generated using NEAT-seq. **g)** TSS enrichment of ATAC-seq data generated using NEAT-seq.

**Supplementary Fig. 2:**
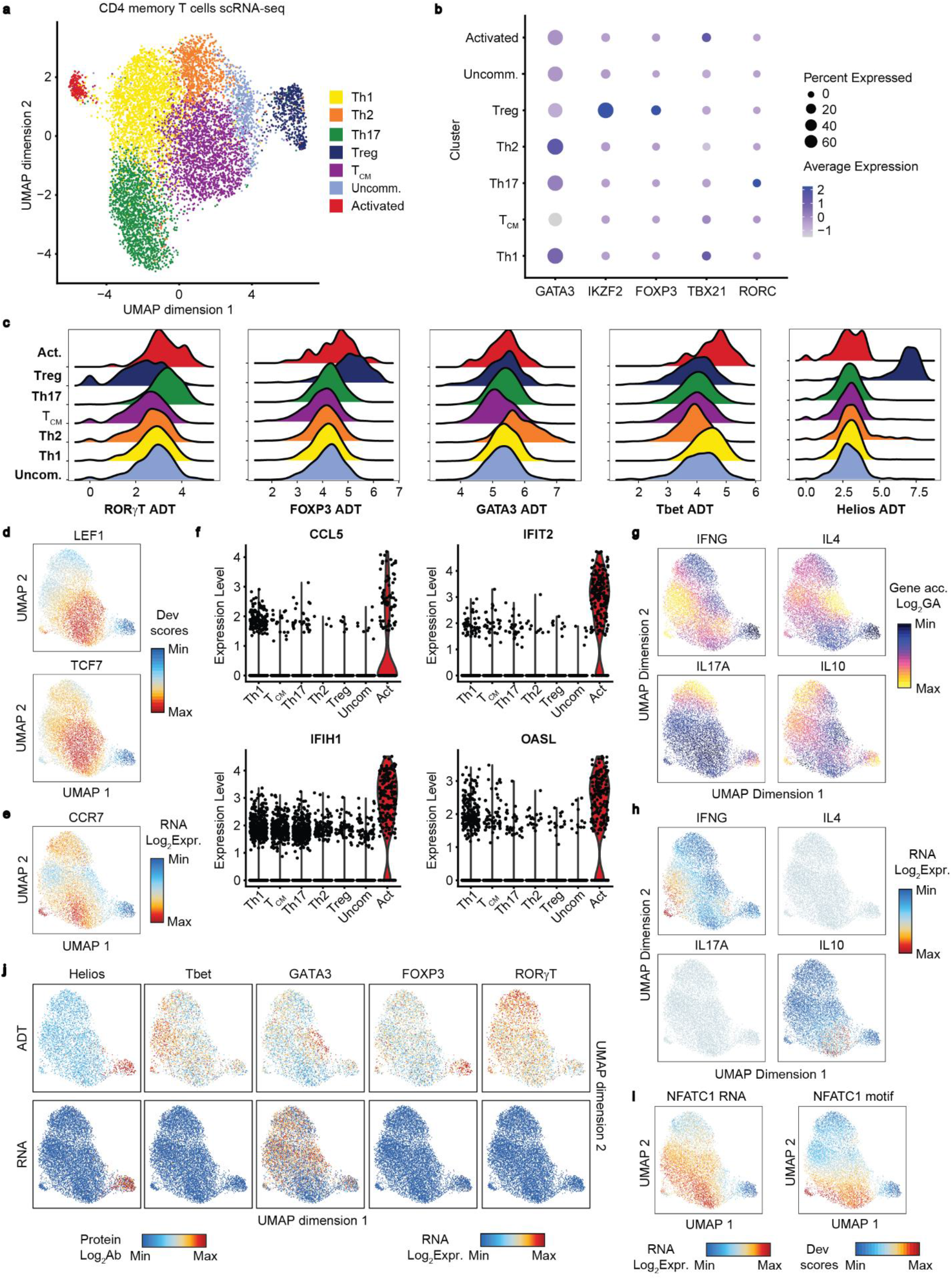
Annotation of scATAC-seq and scRNA-seq clusters. **a)** scRNA-seq UMAP of CD4 memory T cells with cell type classifications. TCM = central memory, Act. = recently activated cells, Uncom. = uncommitted memory cells. **b)** Expression of master TF drivers of CD4 memory cell subsets in each scRNA-seq cluster. TBX21 = Tbet transcript, RORC = RORγT transcript, IKZF2 = Helios transcript. **c)** Log2-transformed, NPC-normalized ADT counts for each TF separated by scATAC-seq cluster**. d)** chromVAR deviation scores for the naïve and CM T cell TFs, LEF1 and TCF7, overlayed on scATAC-seq UMAP. **e)** RNA expression of the CM marker, CCR7, overlayed on the scATAC-seq UMAP. **f)** RNA expression of the indicated genes across clusters in the scRNA-seq UMAP. **g)** Gene accessibility for cytokines induced in different CD4 T cell subsets overlayed on the scATAC-UMAP. **h)** RNA levels for the cytokines in **(g)**. **i)** NFATC1 RNA expression and chromVAR deviation scores overlayed on scATAC-seq UMAP. **j)** Overlay of unsmoothed ADT and RNA levels on the scATAC-seq UMAP.

**Supplementary Fig. 3:**
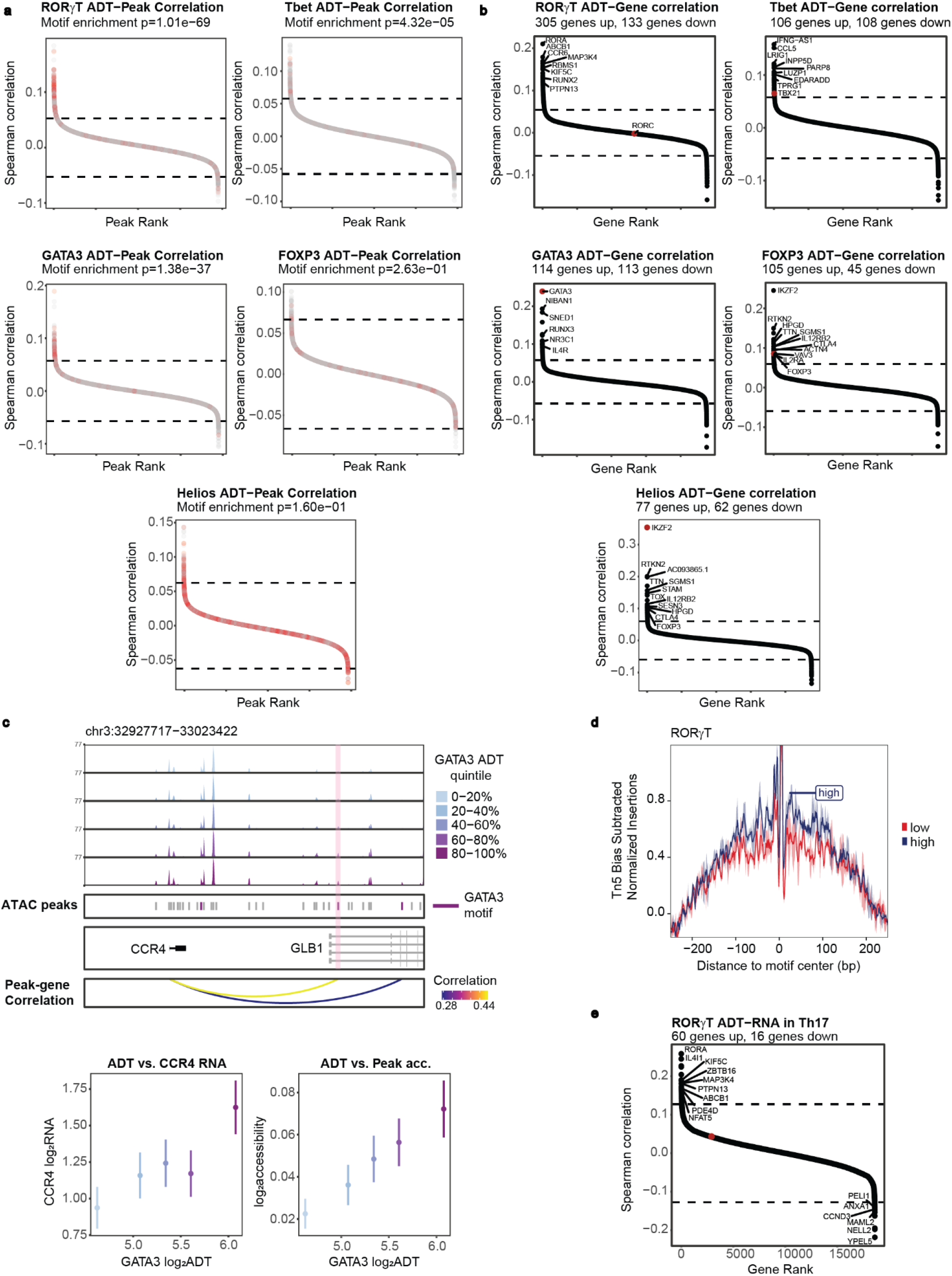
Identification of DNA regulatory elements and genes correlated with master TF protein expression. **a)** Spearman correlations between ATAC-seq peak accessibility and NPC-normalized TF ADT counts across single cells. Cutoffs for significant correlations are indicated by dashed lines (see Methods). Points in red indicate peaks containing a binding motif for the TF. **b)** Spearman correlations between read-normalized RNA counts and NPC-normalized TF ADT counts across single cells. Cutoffs for significant correlations are indicated by dashed lines (see Methods). Significantly correlated genes known to be enriched or play a functional role in the relevant T cell subset are labeled. **c)** Top: CCR4 ATAC-seq tracks in CD4 memory cells separated into quintiles by GATA3 ADT levels, along with significant peak-gene linkages (adj. p < 0.05). Peaks containing a GATA3 motif are indicated. Bottom: CCR4 RNA expression and accessibility at the highlighted GATA3 motif-containing peaks as a function of GATA3 ADT levels. Mean is shown with standard error of the mean. **d)** TF footprinting in ATAC-seq peaks containing the RORγT motif for cells within the annotated Th17 cluster having the highest vs lowest 30% of RORγT ADT levels. ADT counts were NPC-normalized and smoothed across cells with similar ATAC-seq profiles using MAGIC. **e)** Spearman correlations between read-normalized RNA counts and NPC-normalized, smoothed RORγT ADT counts across single cells in the Th17 cluster. Cutoffs for significant correlations are indicated by dashed lines (see Methods). Significantly correlated genes known to be enriched or play a functional role in Th17 cells are labeled. Top anti-correlated genes are also labeled.

